# The mevalonate pathway contributes to monoterpene production in peppermint

**DOI:** 10.1101/2020.05.29.124016

**Authors:** Somnath Koley, Eva Grafahrend-Belau, Manish L. Raorane, Björn H. Junker

**Author notes:** **Authors’ information: Somnath Koley**, **Dr. Eva Grafahrend-Belau**, **Dr. Manish L. Raorane**, **Prof. Dr. Björn H. Junker**. **Materials Distribution:** The author responsible for distribution of materials integral to the findings presented in this article in accordance with the policy described in the Instructions for Authors (www.plantcell.org) is: Björn H. Junker.

## Abstract

Peppermint produces monoterpenes which are of great commercial value in different traditional and modern pharmaceutical and cosmetic industries. In the classical view, monoterpenes are synthesized via the plastidic 2-C-methyl-D-erythritol 4-phosphate (MEP) pathway, while the cytosolic mevalonate (MVA) pathway produces sesquiterpenes. Interactions between both pathways have been documented in several other plant species, however, a quantitative understanding of the metabolic network involved in monoterpene biosynthesis is still lacking. Isotopic tracer analysis, steady state ^13^C metabolic flux analysis (MFA) and pathway inhibition studies were applied in this study to quantify metabolic fluxes of primary and isoprenoid metabolism of peppermint glandular trichomes (GT). Our results offer new insights into peppermint GT metabolism by confirming and quantifying the crosstalk between the two isoprenoid pathways towards monoterpene biosynthesis. In addition, a quantitative description of precursor pathways involved in isoprenoid metabolism is given. While glycolysis was shown to provide precursors for the MVA pathway, the oxidative bypass of glycolysis fueled the MEP pathway, indicating prominent roles for the oxidative branch of the pentose phosphate pathway and RuBisCO. This study reveals the potential of ^13^C-MFA to ascertain previously unquantified metabolic routes of the trichomes and thus advancing insights on metabolic engineering of this organ.

## INTRODUCTION

Essential oils, mainly consisting of volatile terpenes, have huge industrial value due to their commercial uses in aromatic, fragrance, cosmetic, nutraceutical and therapeutic purposes (Duke et al., 2000; Schilmiller et al., 2008). Plants have two independent pathways for supplying precursors of terpene biosynthesis (Figure 1) (Rohmer, 1999). The plastidic 2-C-methyl-D-erythritol 4-phosphate (MEP) pathway begins with condensation of glyceraldehyde 3-phosphate (GAP) and pyruvate (PYR), whereas the cytosolic mevalonate (MVA) pathway is initiated with three molecules of acetyl-CoA (AcCoA). Although both pathways are compartmentally separated inside the plant cell, they synthesize common isomeric isoprenoid units (C_5_), namely isopentenyl pyrophosphate (IPP) and dimethylallyl pyrophosphate (DMAPP). Initial research indicated that the MEP pathway provides the substrates for monoterpenes (C_10_), diterpenes (C_20_) and tetraterpenes (C_40_), whereas the MVA pathway provides the substrates for production of sesquiterpenes (C_15_) and triterpenes (C_30_) (Rodríguez-Concepcíon, 2006). Plant species have evolved several committed organs and cells internally or at the surface, for storage and seclusion of high concentrations of these secondary metabolites (Tissier, 2018). Glandular trichomes (GTs) are one such dedicated extracellular structures, which are found in about 30% of vascular plants (Glas et al., 2012). GTs of the Lamiaceae family exclusively produce and accumulate various volatile terpenes (Lange, 2015).

**Figure 1.**
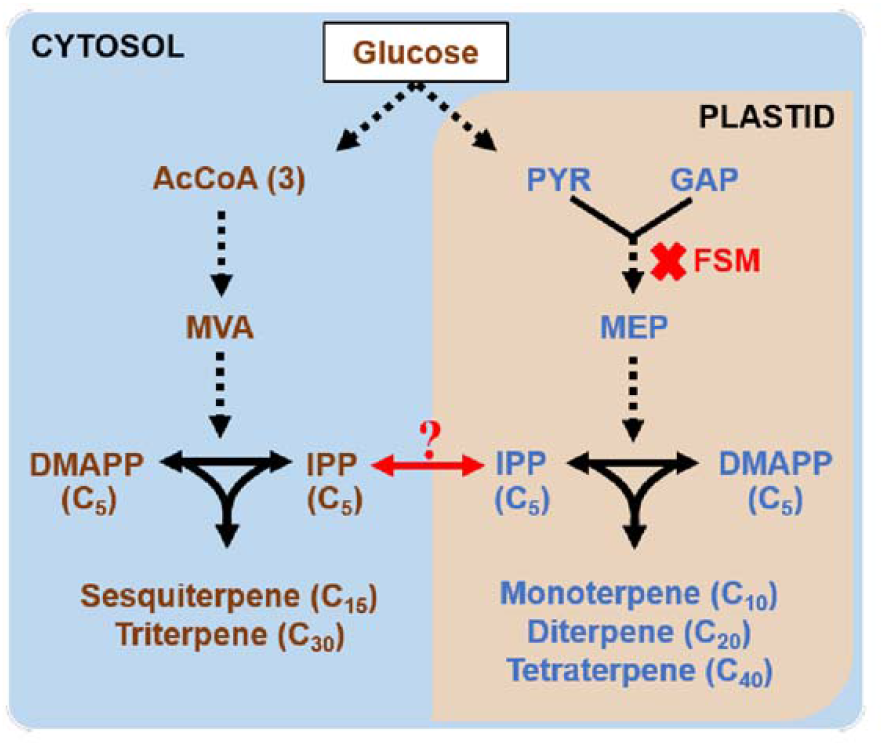
Terpenoid biosynthesis in plants by the means of MEP and MVA pathway. Blue and light orange area indicate the cytosolic and plastidic organelle of the glandular trichome cell, respectively. MEP pathway occurs in plastid and produces mono-, di- and tetraterpene, whereas MVA pathway occurs in cytosol and produces sesqui- and triterpene. Fosmidomycin (FSM) inhibits a key enzyme (DXP reductoisomerase) in the MEP pathway of terpenoid biosynthesis. All abbreviations are listed in Table S7.

Peppermint (*Mentha piperita*), a herb from the Lamiaceae family, is commercially important for its strong aromatic essential oil (Morris, 2006). The principal constituent of the peppermint essential oil are monoterpenes (Rohloff, 1999) which are synthesized and accumulated inside the peltate type GTs to about 88% of its total biomass (Gershenzon et al., 1989; McCaskill et al., 1992). Meta-analysis of peppermint GTs also exhibited a higher degree of metabolic specialization (55% of all GT transcripts) towards terpene biosynthesis (Zager and Lange, 2018). Due to their high biosynthetic activity, these specialized organs of peppermint are extensively used as a model system for terpene biosynthesis during the last three decades.

In the classical view, the MEP pathway has an exclusive role for monoterpene biosynthesis in peppermint GTs (Eisenreich et al., 1997; McCaskill and Croteau, 1995). However, growing evidences of crosstalk between the two isoprenoid pathways have been reported in different plants; such as contribution of the MEP pathway to sesquiterpene production of chamomile (Adam et al., 1999), snapdragon (Dudareva et al., 2005) and cotton (Opitz et al., 2014). This interaction for sterol biosynthesis was evidenced in tobacco bright yellow-2 cells (Hemmerlin et al., 2003) and in arabidopsis (Kasahara et al., 2002). Likewise, the MVA pathway has also been reported for the production of monoterpene (Bartram et al., 2006; Hampel et al., 2007; Mendoza-Poudereux et al., 2015; Opitz et al., 2014; Piel et al., 1998; Schuhr et al., 2003).

Various approaches of ^13^C and ^14^C tracer analysis have been used to understand the interaction between the MEP and MVA pathways (Opitz et al., 2014 and references therein). To study relative pathway contributions, researchers fed various labeled precursors in the form of either general metabolic precursors, such as glucose (GLC) and PYR (Rohmer, 1999; Schuhr et al., 2003), or pathway-specific precursors such as mevalonate and 1-deoxy-D-xylulose (Opitz et al., 2014). Feeding of pathway-specific precursors might alter the physiological condition in the plant by increasing the pool size of the precursor metabolite. Consequently, the respective downstream pathway would be upregulated which finally causes the alteration of the flux split between the two isoprenoid pathways. Opitz et al. (2014) reasoned this as one of the important rationale for the considerable variation in isoprenoid labeling obtained in different tracer studies (Arigoni et al., 1997; Bartram et al., 2006; Dudareva et al., 2005; Hampel et al., 2007; Hemmerlin et al., 2003; Kasahara et al., 2002; Piel et al., 1998; Schuhr et al., 2003; Schwender et al., 1997). Contrastingly, general metabolic precursors such as GLC, do not alter the isoprenoid flux ratio as they exert similar effects on both pathways.

Pathway-specific inhibitor studies (Dudareva et al., 2005; Hemmerlin et al., 2012; Laule et al., 2003 and references therein) and the analysis of transgenic plants (e.g., Carretero-Paulet et al., 2006; Neelakandan et al., 2010; Rodríguez-Concepción et al., 2004) have also been used to quantify relative pathway usage. These approaches may have off-target effects and perturb the cellular system (Shih and Morgan, 2020). For example, Laule et al. (2003) reported a complementation effect exhibited by one pathway due to the inhibition of the other. Thus, inhibition and transgenic studies should only be used as an additional support for flux quantifications that have been carried out under conditions that are not perturbing the pathways.

Although all these studies indicate crosstalk between the MEP and MVA pathways as a common phenomenon in plants, a quantitative understanding of how isoprenoid and associated primary metabolic pathways are contributing to isoprenoid production is still lacking. While the above-mentioned methods are restricted to quantify only the split ratio between the two isoprenoid pathways, a system-wide quantification of isoprenoid and primary metabolism is essential to be able to efficiently engineer isoprenoid biosynthesis.

Metabolic flux analysis (MFA) is a method to quantify intracellular fluxes at the network level (Ratcliffe and Shachar-Hill, 2005). The resulting flux maps provide a quantitative measure of the metabolic phenotype as they reflect the integrated phenotype of genome, transcriptome, proteome and regulatory interactions (Stephanopoulos, 1999). In the last two decades, different MFA approaches have been applied to plant systems (for a review, see Kruger et al., 2012; Kruger and Ratcliffe, 2015), with steady state ^13^C-MFA being the gold standard method (Ratcliffe and Shachar-Hill, 2006; Schmidt et al., 1997; Wiechert, 2001; Zupke and Stephanopoulos, 1994). Steady state flux analysis has proved to be a valuable tool to investigate the effect of environmental, and genetic changes (Alonso et al., 2007; Junker et al., 2007; Masakapalli et al., 2014, 2013; Rontein et al., 2002; Williams et al., 2008). It has also resolved the fluxes of parallel metabolic routes (Allen et al., 2009; Stephanopoulos, 1999) and identified novel alternate pathways (Schwender et al., 2004).

In plant science, steady state MFA has been mainly focused on primary carbon metabolism, while labor- and time-intensive kinetic modeling or dynamic labeling has been used to study fluxes in secondary metabolism (Boatright et al., 2004; Matsuda et al., 2005, 2003; Okazaki et al., 2004; Orlova et al., 2006; Rios-Estepa et al., 2008; Zhang et al., 2015). However, given that the basic requirement for steady state MFA are fulfilled (e.g. mixo- or heterotrophic conditions, rearrangement of the labeled core carbon structure), this technique can also be applied to secondary metabolism. It allows resolving parallel pathways under physiological conditions, quantifying primary metabolism around key intermediary precursors of secondary metabolites and detecting fluxes to wasteful competitive pathways (Shih and Morgan, 2020); thus, steady state MFA becomes a powerful method for investigating plant secondary metabolism in a system-wide and quantitative manner.

To study the contribution of the MEP and MVA pathway to monoterpene biosynthesis in peppermint GT, we performed ^13^C tracer analysis and pathway inhibition studies. Additionally, steady state ^13^C MFA using data sets of three different tracer studies was applied to quantify fluxes of the two isoprenoid pathways and their precursor pathways of primary metabolism. Our approach offers improved insights into peppermint GT metabolism by (1) providing evidence for the contribution of the alternate MVA pathway to monoterpene biosynthesis and (2) quantifying the crosstalk between the MEP and MVA pathway along with primary metabolism involved in monoterpene production of peppermint GT. This study reveals the potential of steady state ^13^C-MFA to ascertain previously unquantified metabolic routes within the trichome cell and thus providing assured metabolic engineering targets aimed towards increased production of these high value compounds.

## RESULTS

To quantify fluxes of primary and secondary metabolism involved in monoterpene production in non-photosynthetic GTs of peppermint - ^13^C tracer analysis, pathway inhibition studies and steady state ^13^C-MFA were performed using isotopic labeling data of three different tracers (1-^13^C_1_, 6-^13^C_1_ and 1,6-^13^C_2_ GLC).

### Metabolic and isotopic steady state

For the application of steady state ^13^C tracer analysis and ^13^C-MFA, the in vitro system has to be in metabolic and isotopic steady state (Roscher et al., 2000). In the present study, the production of monoterpenes was analyzed in the leaves of the peppermint plants generated through earlier established shoot-tip culture method (Koley et al. 2019). Due to their trichome-specific biosynthesis, volatile monoterpenes are well suited to quantify GT metabolism without the need for trichome isolation. The linear accumulation phase of monoterpene was observed between 13 and 16 days (Figure 2A), indicating metabolic steady production of monoterpene in peppermint GTs. Isotopic enrichment in monoterpene was stable during 14 to 16 days age of the first leaf-pair (Figure 2B) with an average label incorporation of 99.5% in 15 days old leaf-pair grown in U-^13^C_6_ GLC media. Both, stability and proximity to the maximum possible enrichment (100%) indicate isotopically unchanged conditions, thus confirming an isotopic steady state at the end of the cultivation period in GTs. Therefore, further studies were performed with 15 days aged leaf-pairs of these peppermint plants.

**Figure 2.**
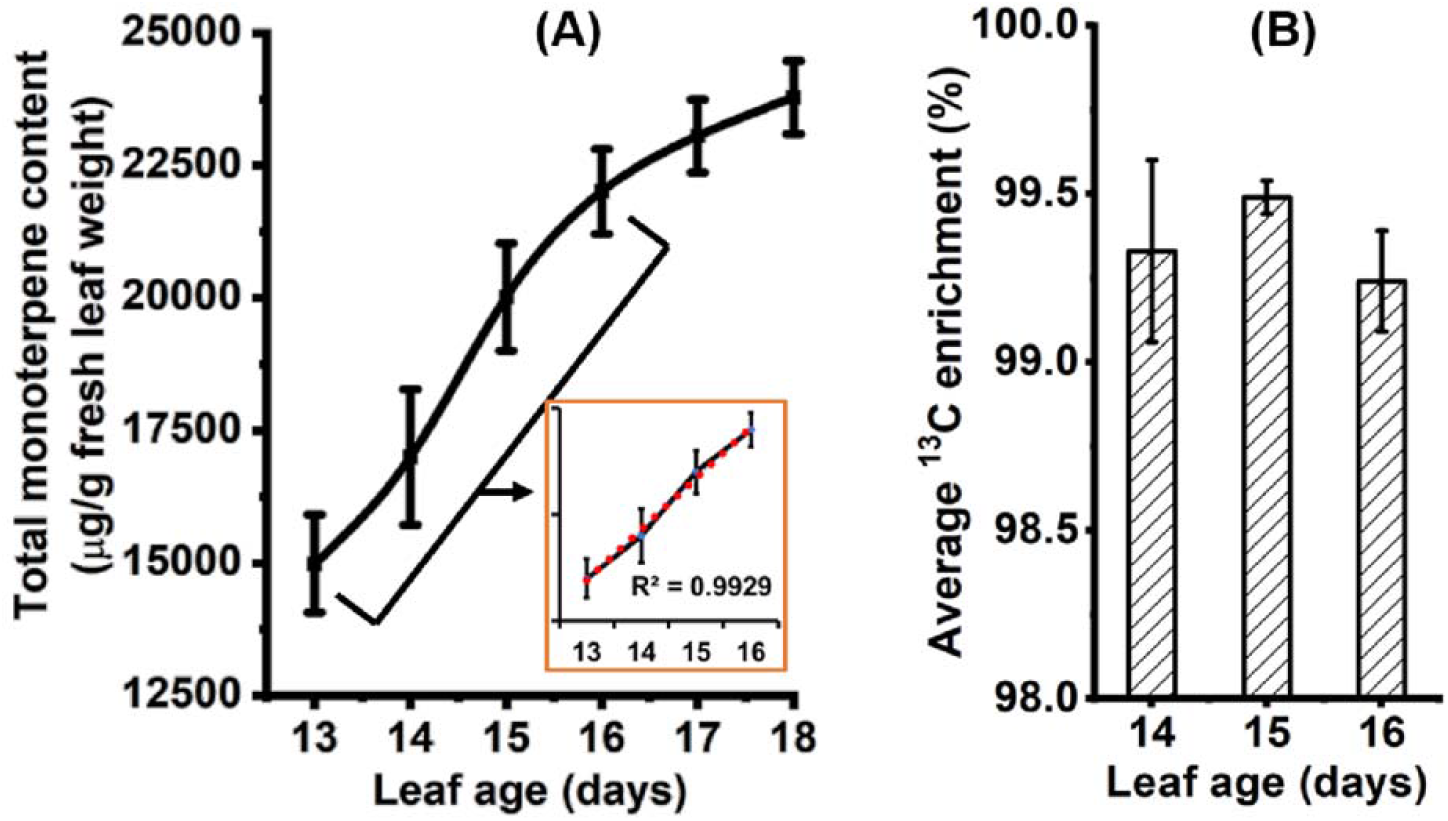
Steady state determination in the first leaf pair of peppermint shoot-tip culture. **(A)** Determination of metabolic steady state by quantifying total monoterpene where steady production of the monoterpene in vitro culture was observed during 13 to 16 days. R-squared value of liner regression trendline for that growth period is zoomed to depict the steady state. **(B)** Determination of isotopic steady state by assessing average ^13^C enrichment. The culture was grown under 5 μmol m^−2^s^−1^ light intensity in the basal media containing **(A)** unlabeled and **(B)** U-^13^C_6_ glucose (mean ± SE, n=5).

### Isotopic tracer analysis

Three different isotopic GLC tracer studies were performed to decipher GT metabolism. As shown by the ^13^C labeling analysis of monoterpene, the mass isotopomer distribution (MID) of pulegone differed between all three tracer studies (Figure 3). Labeling with [1-^13^C_1_]-GLC resulted in a significant degree of M+0 and M+1 pulegone, while a higher percentage of M+2 labeled pulegone was observed from [6-^13^C_1_]-GLC source. Interestingly labeling with [1,6-^13^C_2_]-GLC resulted in a higher degree of M+3 pulegone and the emergence of M+5 and M+6 labeled pulegone. Isotopic patterns similar to pulegone, were also confirmed by two other monoterpenes (menthofuran and isopulegone, Table S2 and S3).

**Figure 3.**
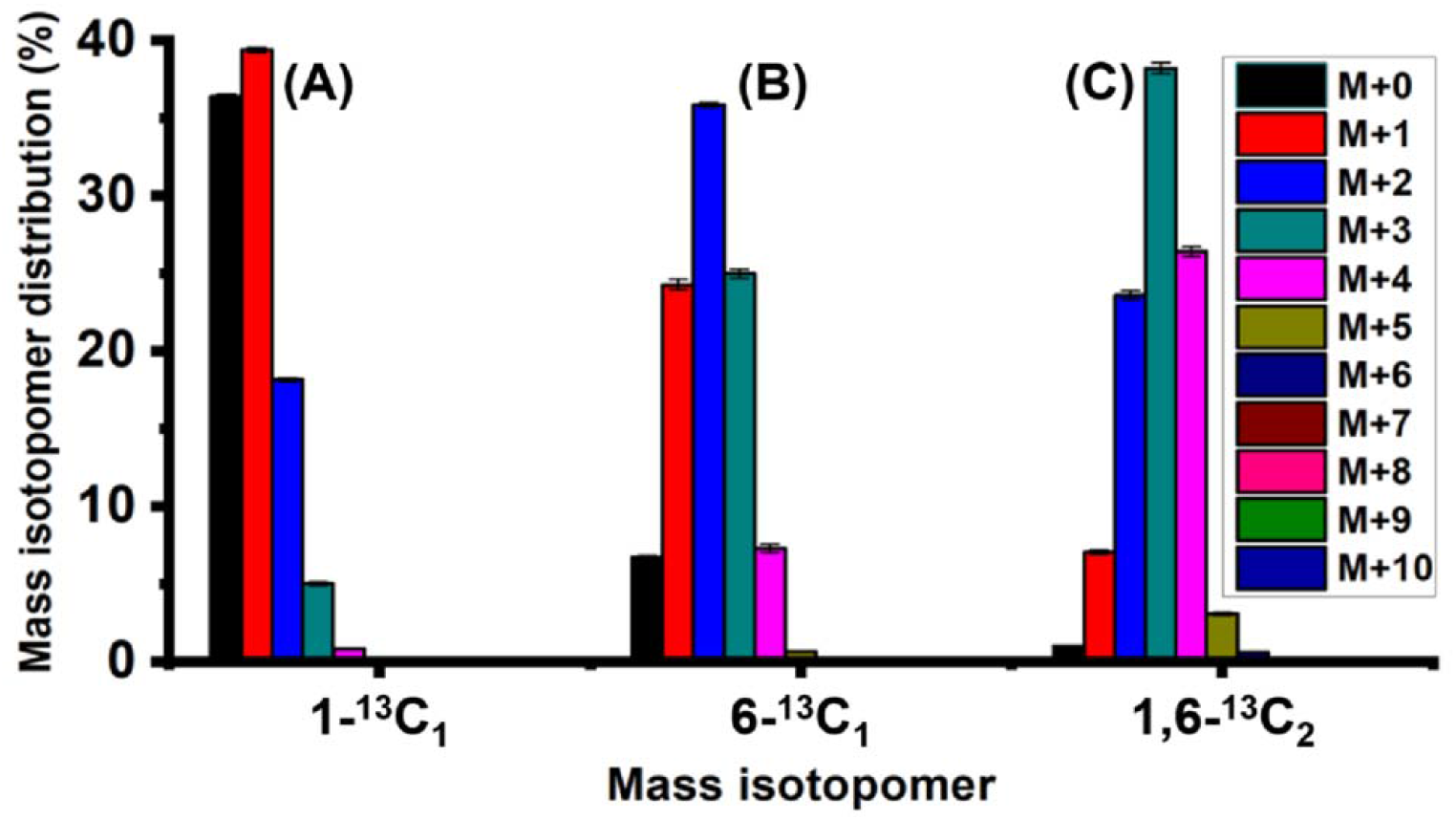
Mass isotopomer distribution of pulegone measured by GC-MS In the first leaf pair for three different tracer experiments. Peppermint shoot-tip culture was grown under 5 μmol m^−2^ s^−1^ light intensity in the basal media containing **(A)** 1 -^13^C_1_, **(B)** 6-^13^C_1_ and **(C)** 1,6-^13^C_2_ glucose as the sole carbon source (mean ±SE, n=3). Similar MID patterns were also found in other monoterpenes and present in Table S2 and Table S3.

As depicted in the theoretical scheme of carbon atom transitions from three applied tracers (Figure 4), M+0 to M+4 labeled monoterpenes can potentially be formed by the MEP and MVA pathway, whereas M+5 and M+6 monoterpenes production is specific to the MVA pathway. Hence, a significant amount of M+5 and M+6 monoterpene measured in the [1,6-^13^C_2_]-GLC tracer study indicated the activity of the cytosolic MVA pathway for monoterpene production.

**Figure 4.**
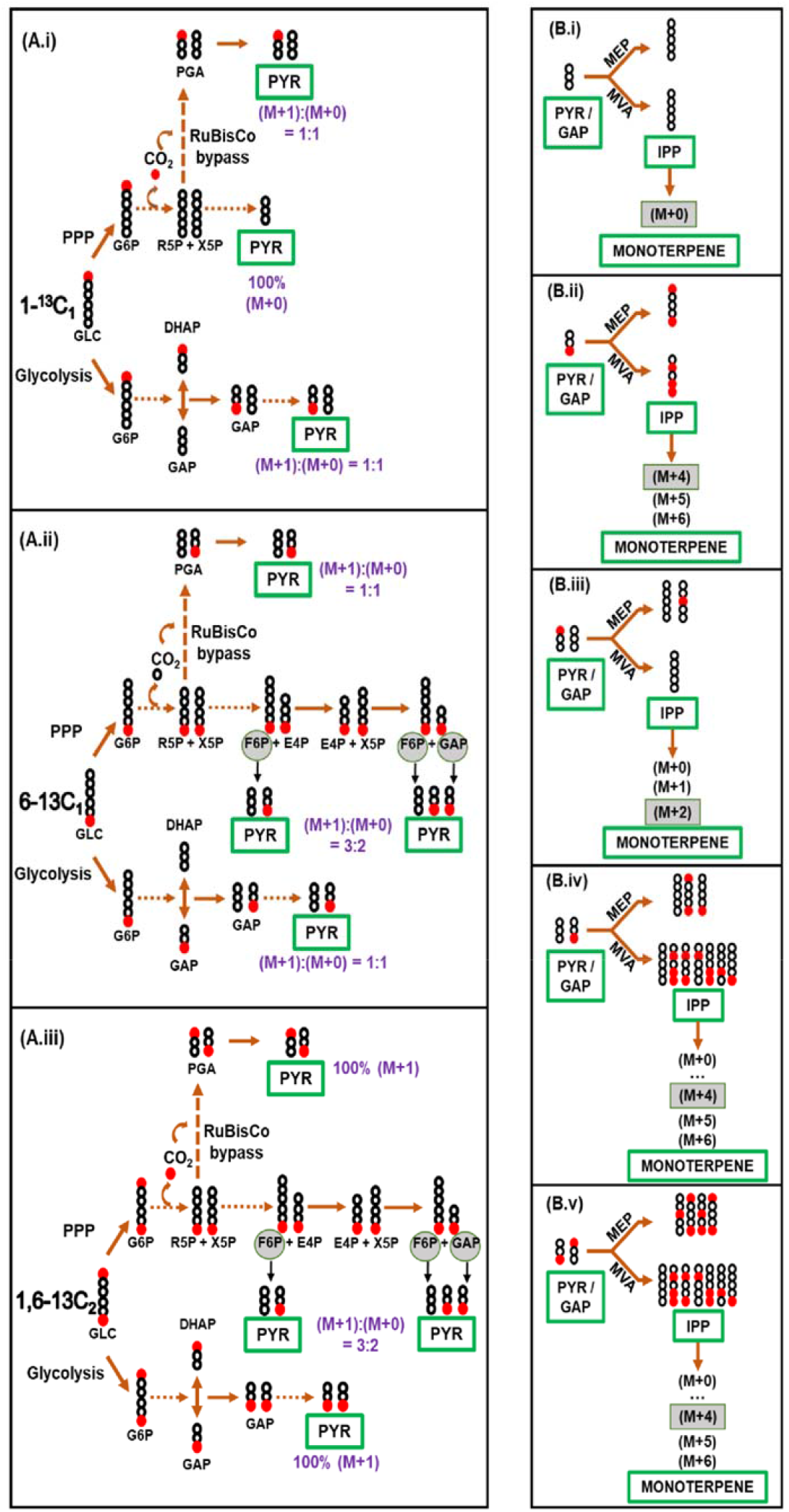
Theoretical overview of ^13^C labeling patterns derived from three different GLC tracers. Simplified scheme of ^13^C label derived from 1-^13^C_1_, 6-^13^C_1_ and 1,6-^13^C_2_ glucose via **(A. i-iii)** glycolysis, PPP and RuBisCO. Source of trioses from PPP is encircled with grey background for visual understanding. PYR and GAP produced from the above-mentioned three pathways are metabolized by the **(B. i-v)** MEP and MVA pathway for monoterpene production. In MEP pathway, second and third carbon of PYR are attached to the whole GAP molecule to produce C_5_ unit; first carbon of PYR is released as CO_2_. In MVA pathway, first carbon of three PYR molecule was released for the production of three AcCoA, from where another carbon (equivalent to second carbon of PYR) of one random AcCoA was released for the production C_5_ unit. In this comprehensive analysis, any of the two C_5_. units regardless of the pathway are expected to be joined together to produce one monoterpene. The highest possible mass isotopomer of monoterpene, synthesized exclusively via the classical MEP pathway, is marked by a grey rectangular box in B.i-v. Unlabeled carbon (^12^C) and isotopic carbon (^13^C) are denoted by black and red circle. All abbreviations are listed in Table S7.

The ^13^C labeling data also suggested an elevated flux of oxidative pentose phosphate pathway (oxPPP) in non-photosynthetic GTs. As depicted in Figure 4 (A.i-A.ii), glycolytic pathway alone would not lead to any difference in labeling pattern between [1-^13^C_1_]- and [6-^13^C_1_]-GLC fed cultures. The RuBisCO shunt (i.e. the combination of reverse non-oxPPP and subsequent CO_2_ fixation by RuBisCO activity) would also produce the identical labeling phenotype (Figure S2). However, higher differences were observed in M+0 monoterpene of these two tracer studies (Figure 3), denoting an increased metabolism of oxPPP, instead of sole glycolytic route (Figure 4A.i-A.ii, and 4B.i-B.iv).

Isotopic analogy in monoterpenes from [6-^13^C_1_]- and [1,6-^13^C_2_]-GLC tracer experiments implied RuBisCO to play a role in peppermint GT metabolism. The distribution of M+0 to M+3 labeled monoterpene from [6-^13^C_1_]-GLC was roughly equal to M+1 to M+4 labeled monoterpene from [1,6-^13^C_2_]-GLC (Figure 3). The observed one-shift MID pattern can only result from the conversion of the oxPPP-derived pentose phosphate by the route of RuBisCO fixation. While the same amount of isotopic carbon is passed via non-oxPPP using [6-^13^C_1_]-and [1,6-^13^C_2_]-GLC, one extra ^13^C from [1,6-^13^C_2_]-GLC is provided via the RuBisCO bypass (Figure 4A.ii and 4A.iii). In the next step, PYR from [1,6-^13^C_2_]-GLC produces more ^13^C enriched IPP via MEP pathway, compared to [6-^13^C_1_]-GLC (Figure 4B.iv and 4B.v). Neither glycolysis nor the combination of glycolysis and PPP without RuBisCO fixation can eventuate the feature of one-shift MIDs from M+1 to M+4, as the involvement of glycolysis can only increase M+4 enrichment of monoterpene from [1,6-^13^C_2_]-GLC, contrasting from [6-^13^C_1_]-GLC (Figure 4A.iii and 4B.ii). Additionally, very little M+0 labeled monoterpene from the experiment of [1,6-^13^C_2_]-GLC contradicted the presence of significant non-oxPPP pathway (Figure 3C, and Figure 4A.iii and 4B.iv).

### Inhibition study

Inhibition studies were performed to confirm the alternate contribution of the MVA pathway to monoterpene biosynthesis as predicted by ^13^C tracer analysis. As a potent inhibitor of the MEP pathway, fosmidomycin (FSM) was applied to study the contribution of the MVA pathway to the biosynthesis of classical MEP pathway-derived compounds such as carotenoids and phytol side chains of chlorophylls. Four different concentrations of FSM were tested to observe the effect of inhibition on total chlorophyll and carotenoid content. Although comparable growth of the shoot tip culture was observed, phenotypic difference between control and treated plants indicated the inhibition of chlorophyll biosynthesis (Figure S1). Inhibition for phytopigment production was observed for treatment with 20, 30 and 40 μM FSM (Table 1). Nevertheless, a relatively small quantity of carotenoids and chlorophylls were still produced, even after the highest level of MEP inhibition, thus suggesting a plausible contribution of the alternate MVA pathway in peppermint leaf metabolism.

**Table 1.** Production of phytopigments from fosmidomycin treated shoot-tips. Fifteen days old leaf pairs were obtained from growing shoot-tips cultivated in culture media containing 0 (control), 10, 20, 30 or 40 μM FSM (mean ± standard error, n=5).

| Fosmidomycin concentration | Phytopigments content (mg/g fresh weight) |  |
| --- | --- | --- |
|  | Total chlorophyll | Total carotenoid |
| 0 $\mu\text{M}$ | 1.75 ± 0.19 | 0.25 ± 0.02 |
| 10 $\mu\text{M}$ | 0.20 ± 0.02 | 0.06 ± 0.01 |
| 20 $\mu\text{M}$ | 0.06 ± 0.02 | 0.02 ± 0.01 |
| 30 $\mu\text{M}$ | 0.05 ± 0.02 | 0.02 ± 0.01 |
| 40 $\mu\text{M}$ | 0.07 ± 0.01 | 0.02 ± 0.01 |

Although the production of the photosynthetic pigments were only partially blocked at 10 μM FSM, this concentration of the inhibitor was deemed ideal for further ^13^C based studies as a certain quantity of metabolite is necessary for precise analysis of MIDs. This concentration of FSM was added to the medium in a [1-^13^C_1_]-GLC tracer experiment to comprehend the contribution of MVA pathway based on the rationale that similar isotopic profile in pulegone of control vs. partially-inhibited GTs indicates monoterpene biosynthesis via the classical MEP pathway, whereas any observed differences indicate the role of the alternate MVA pathway in monoterpene production. As shown in Figure 5, the average ^13^C enrichment in monoterpene was higher in inhibited GTs (13%) compared to control (9.5%). In addition, the lower mass isotopomers, such as M+0 and M+1 pulegone were reduced after inhibition, whereas M+2 to M+5 pulegone were increased, compared to the control. Theoretical depiction of [1-^13^C_1_]-GLC flow demonstrates the presence of extra isotopic carbon via the MVA pathway, as this pathway involves three trioses as compared to two trioses in case of the MEP pathway (Figure S3). Likewise, in the present study, a consequence of the MEP pathway inhibition by FSM led to the elevation in average ^13^C enrichment and distribution of higher mass isotopomers; strongly indicating the role of the MVA pathway to monoterpene production in peppermint GT.

**Figure 5.**
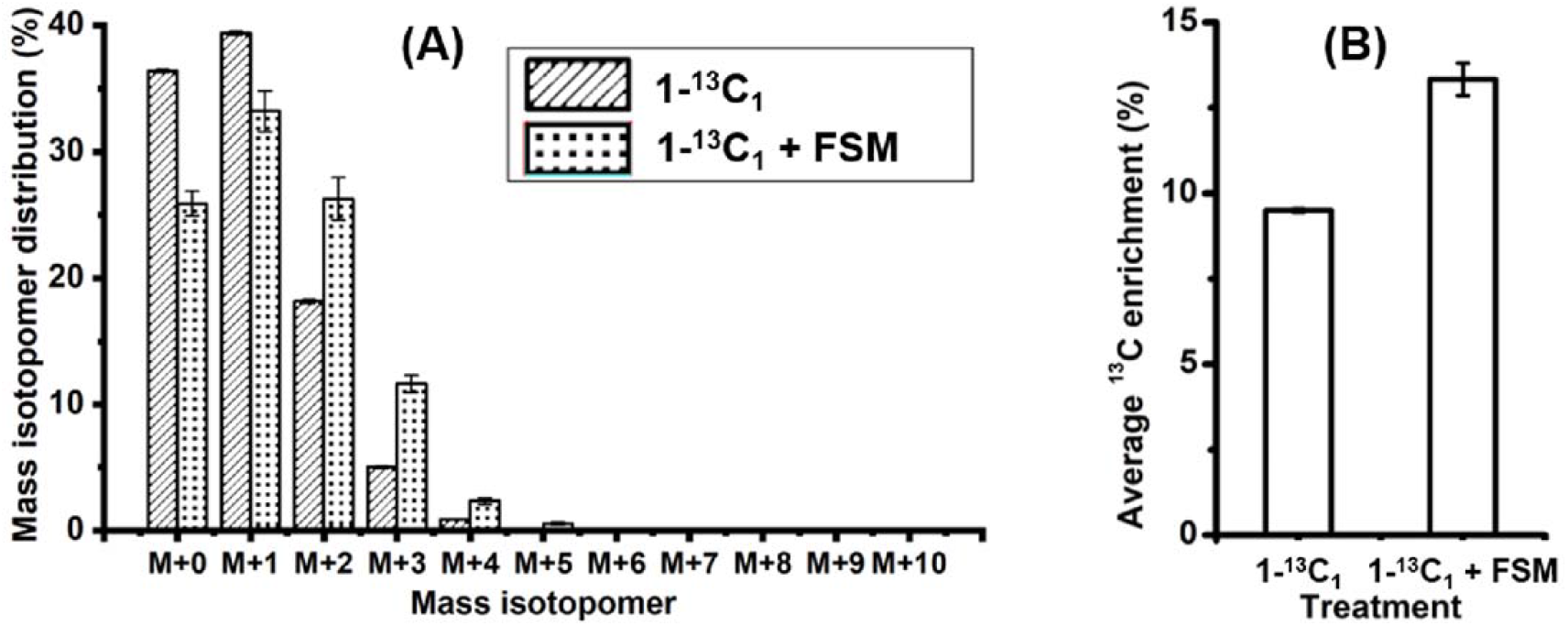
Labeling study after partial inhibition of the MEP pathway. Fifteen days old leaf pairs were obtained from growing shoot-tips cultivated in media containing either 1 -^13^C_1_ glucose or with 1-^13^C_1_ glucose + FSM (FSM-10 μM concentration). **(A)** MID and **(B)** average ^13^C enrichment of pulegone were assessed from the control and inhibited culture (mean ± SE, n=3).

### Steady state ^13^C-MFA

The findings described above, point to a role for the alternate MVA pathway in monoterpene production by peppermint GTs. In addition, ^13^C tracer analysis suggested the oxPPP and RuBisCO to be an integral part of central carbon metabolism in non-photosynthetic GTs. To fully resolve and quantify the metabolic fluxes involved in GT metabolism, we performed parallel steady state ^13^C-MFA. Isotopic labeling data from monoterpenes of the above-described tracer studies (TableS1, S2 and S3) was simultaneously fitted to a compartmentalized model of monoterpene biosynthesis within non-photosynthetic peppermint GTs during secretory phase. In the step-wise process of metabolic model reconstruction, first central pathways involved in monoterpene biosynthesis were identified by surveying the respective biochemical literature (e.g., (Ahkami et al., 2015; Gershenzon et al., 1989; Johnson et al., 2017)) and online databases of Brenda (Placzek et al., 2017), MetaCyc (Caspi et al., 2018) and PlantCyc (Dreher, 2014). In the next step, the initial model was refined in an iterative data-driven process based on a comparative analysis of alternative model versions. Variations tested in the different models comprise (1) in-/exclusion of certain reactions/pathways (e.g. fermentation, cytosolic oxidative pentose phosphate pathway), (2) different degrees of compartmentalization (e.g. isoenzymes, combined metabolite pools) and (3) model simplifications (e.g. lumping of linear pathways).

The final model used for ^13^C-MFA comprised 37 biochemical reactions and 12 transport processes across four different compartments (i.e. cytosol, mitochondria, leucoplast, extracellular medium; Table S4). In addition to monoterpene biosynthesis (MEP and MVA pathway), the compartmentalized model integrated glycolysis, the tricarboxylic acid (TCA) cycle, the PPP, the RuBisCO bypass (Schwender et al., 2004) and multiple ana-/cataplerotic reactions. Imported GLC is converted to precursors for monoterpene biosynthesis via glycolysis (cytosolic and plastidic), and the oxPPP and non-oxPPP. The resulting triose phosphates, namely, plastidic GAP and PYR serve as precursors for the MEP pathway. Alternatively, plastidic PYR can also be synthesized via the plastidic NADP-dependent malic enzyme. Cytosolic AcCoA, the precursor for the MVA pathway, is obtained via the citrate (CIT) shuttle. In the mitochondria, CIT is produced by citrate synthase via the TCA cycle, and then transported into the cytosol where it is cleaved to oxaloacetate and AcCoA by the cytosolic ATP-citrate lyase. To replenish the carbon withdrawn from the TCA cycle, malate (MAL) and PYR are produced via the anaplerotic reaction of phosphoenolpyruvate carboxylase (PEPC), malate dehydrogenase (MDH), malic enzyme (ME) and pyruvate kinase (PK) in the cytosol. Additionally, RuBisCO which was shown to be highly expressed in peppermint GTs (Ahkami et al., 2015), acts as a potential mechanism to recycle CO_2_. To account for the energy and redox-related processes involved in monoterpene biosynthesis, the model also integrates ATP, NADH and NADPH production and consumption. Oxidative phosphorylation is defined based on the stoichiometries of 2.5 times ATP per NADPH molecule (Hinkle, 2005).

To reduce model complexity, the subcellular compartmentation of certain metabolic processes was simplified. As preliminary model versions demonstrated that (1) metabolic fluxes of upper glycolysis could not be precisely resolved based on the provided labeling data, and (2) median flux values of the compartmented vs. uncompartmented flux model did not differ and hence, the compartmentation of the upper part of glycolysis (G6P to GAP) was not considered in the final model. In addition, monoterpene biosynthesis was confined to the plastid, as the first step of monoterpene biosynthesis (from geranyl pyrophosphate to limonene) exclusively takes place in plastid (Alonso et al., 1992; Colby et al., 1993). For further reduction of the model complexity, the exchange of intermediates between the cytosol and mitochondria were restricted to PYR, MAL and CIT, and between cytosol and plastid to G6P, F6P, GAP, PYR, MAL and IPP. The inclusion of alternate triose phosphate and hexose phosphate transporters was tested in preliminary studies and rejected due to their negligible impact on the overall carbon flux. The C_5_ unit or C_5_ derivative of this unit was reported to be transported between the cytosol and plastid (Bick and Lange 2003). Thus, IPP which is the first common intermediate of cytosolic and plastidic terpene biosynthetic pathways was used as the means of crosstalk between these pathways. To account for the dilution of unlabeled carbon sources, the model integrated uptake of unlabeled GLC. Further details of the peppermint GT model are provided in Table S4.

### Metabolic fluxes in non-photosynthetic peppermint GTs

The parallel ^13^C-MFA provided a statistically acceptable fit with the resulting minimized SSR of 79.2 being in the range of the expected lower (40.5) and upper bound (83.3) of the 95% confidence region of SSR. The metabolic flux map representing the best-fit solution is given in Figure 6. The simulated fluxes and the lower and upper bound of the 95% confidence intervals for all fluxes are listed in Table S6. The most important relative fluxes are presented in Table 2 for detailed interpretation of the GT metabolism.

**Figure 6.**
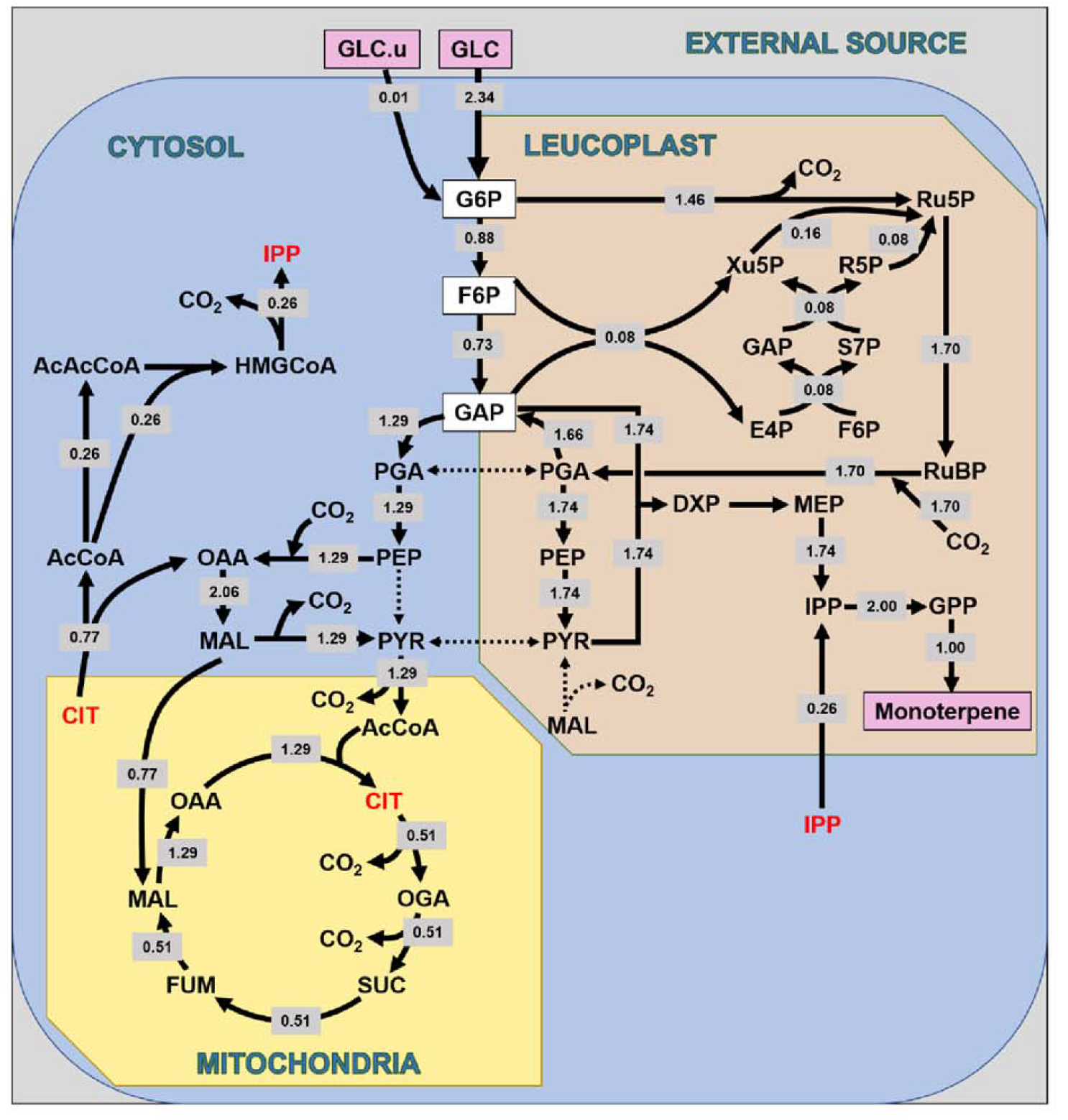
Metabolic flux map of primary and monoterpene metabolism of secretory cells in non-photosynthetic peppermint GTs during the secretory phase. Metabolic fluxes were quantified by parallel ^13^C-MFA using ^13^C labeling patterns of monoterpenes (pulegone, meπthofuran and isopulegone; see Table S1-S3 for replication data sets) derived from 1- ^13^C_1_, 6-^13^C_1_ and 1,6-^13^C_2_ glucose experiments (SEM, n = 3). All estimated fluxes (in grey box) are normalized to a net monoterpene production rate of one. Arrows indicate the direction of net flux, and black dashed arrows indicate zero flux value. Blue, light orange and light yellow area indicate the cytosolic, plastidic and mitochondrial organelle of the glandular trichome cell, respectively. Metabolites in red color (mitochondrial CIT and cytosolic IPP) have been represented twice for visual purposes, although they belong to the same CIT and IPP pool, respectively. All abbreviations used in the figure are listed in Table S7.

**Table 2.** Relative contribution of metabolic fluxes in peppermint GTs quantified by ^13^C-MFA. All estimated fluxes are normalized to a net monoterpene production rate of one. Relative values are enumerated from peppermint GT ^13^C-MFA results. Relative flux in row B and D of the table are predicted from the GAP pool by observing the relative pool of cytosolic and plastidic PGA.

|  | Metabolic fluxes | Total flux | Relative flux | Relative flux (in %) |
| --- | --- | --- | --- | --- |
| A | **Total IPP for monoterpene biosynthesis** | 2 |  |  |
|  | From MEP pathway |  | 1.742 | 87% |
|  | From MVA pathway |  | 0.258 | 13% |
| B | **GAP source for MEP pathway** | 1.742 |  |  |
|  | Produced by glycolysis |  | 0.086 | 5% |
|  | Produced by oxPPP + RuBisCO |  | 1.656 | 95% |
| C | **Glucose conversion** | 2.347 |  |  |
|  | Converted by oxPPP |  | 1.464 | 62% |
|  | Converted by glycolysis |  | 0.726 | 31% |
|  | Converted by non-oxPPP |  | 0.157 | 7% |
| D | **Glycolytic GAP for terpene biosynthesis** | 1.453 (31%) |  |  |
|  | For plastidic MEP pathway |  | 0.165 | 11% (3%) |
|  | For cytosolic MVA pathway |  | 1.288 | 89% (28%) |
| E | **Ru5P source for RuBisCO flux** | 1.699 |  |  |
|  | Produced by oxPPP |  | 1.464 | 86% |
|  | Produced by non-oxPPP |  | 0.235 | 14% |
| F | **Mitochondrial CIT flux** | 1.288 |  |  |
|  | Export to cytosol |  | 0.773 | 60% |
|  | Used for lower TCA cycle |  | 0.515 | 40% |
| G | **Carbon conversion** | 14.082 |  |  |
|  | Carbon converted into biomass |  | 10 | 71% |
|  | Carbon released as CO<sub>2</sub> |  | 4.082 | 29% |
| H | **Total CO<sub>2</sub> production** | 7.069 |  |  |
|  | Total CO<sub>2</sub> fixation |  | 2.987 | 42% |
|  | Total CO<sub>2</sub> release |  | 4.082 | 58% |
| I | **CO<sub>2</sub> fixed by RuBisCO** | 1.699 |  | 24% |
|  | Emerged from oxPPP |  | 1.464 | 21% |
|  | Emerged from other pathway |  | 0.235 | 3% |
| J | **NADPH source** | 7.743 |  |  |
|  | Produced by oxPPP |  | 2.928 | 38% |
|  | Produced by cytosolic NADP-ME |  | 1.288 | 17% |
|  | Consumption from external source |  | 3.527 | 46% |
| K | **NADH source** | 7.564 |  |  |
|  | Produced by glycolysis |  | 1.288 | 17% |
|  | Produced by TCA cycle |  | 3.606 | 48% |
|  | Produced by monoterpene biosynthesis |  | 1 | 13% |
|  | Consumption from external source |  | 1.67 | 22% |
| L | **NADH requirement** | 7.564 |  |  |
|  | Requirement for ATP conversion |  | 3.847 | 51% |
|  | Requirement for other pathways |  | 3.717 | 49% |

#### Crosstalk between MEP and MVA pathway for monoterpene biosynthesis

As shown in Figure 6, the MFA study provided further evidence for a crosstalk between the two isoprenoid pathways. IPP was shown to be produced via the cytosolic MVA pathway and transported into the plastid for the production of geranyl pyrophosphate, the final precursor for monoterpene biosynthesis. Flux analysis indicated that 13% (± 2% SE) of monoterpenes were synthesized via the MVA pathway and 87% (± 2% SE) via the MEP pathway (Table 2A and Table S6). Given a lower bound of the relative MVA flux of 9.5% at 95% CI, the quantified contribution of MVA flux was statistically significant. This is statistically accurate even at a 99.9% confidence level where the interval between 6.9% and 20% contained the true value of the MVA flux for monoterpene biosynthesis (Table S5).

#### Central carbon fluxes towards monoterpene biosynthesis

Precursors of the classical plastidic MEP pathway, namely GAP and PYR, were shown to be predominantly synthesized via the glycolytic bypass of oxPPP and RuBisCO. This oxidative bypass of glycolysis fulfilled the entire PYR demand for MEP pathway, however, a small part (5%) of GAP precursor was exclusively supplied by glycolysis (Table 2B). The plastidic malic enzyme (NADP-ME) carried no flux.

Imported GLC in the secretory cells of peppermint GT was predominantly directed towards the oxidative branch of PPP (62%) (Table 2C), as the major monoterpene biosynthesis pathway (MEP) takes place in the plastid. Besides, GLC was also metabolized by the non-oxPPP (7%) and glycolysis (31%), fueling biosynthesis of triose phosphates. Due to the computed inactivity of transport processes involved in lower glycolysis (by either PYR or PGA), it can be concluded that 11% of total glycolytic GAP (equivalent to 3% of total GLC uptake) was used by plastidic MEP pathway, while cytosolic MVA route utilized 89% (equivalent to 28% of total GLC uptake) (Table 2D). With respect to the upper part of central metabolism in peppermint GT, the relative flux analysis indicated that a minimum of 28% or maximum 31% (when 3% was transported to plastid for DXP pathway) of carbon was metabolized via cytosolic MVA pathway. RuBisCO enzyme was shown to carboxylate Ru5P of oxPPP (86%) and non-oxPPP (14%) to triose phosphates (PGA) (Table 2E). Cytosolic PGA was converted to cytosolic AcCoA (precursor of MVA pathway) via the combined action of TCA cycle and the CIT-MAL shuttle. Fueled by the import of cytosolic PYR and MAL, mitochondrial CIT was exported from the mitochondria to sustain monoterpene biosynthesis (60%). In addition, 40% of CIT was turned into isocitrate to provide energy and redox equivalents via the forward cyclic flux through the lower part of TCA cycle (Table 2F).

#### Carbon conversion efficiency and redox requirements for monoterpene biosynthesis

Metabolic flux analysis indicated that secretory cells of peppermint GTs were highly efficient in converting carbon supplies (GLC) into monoterpenes. For each mol of C_10_ terpene production, cells imported 2.35 mol of C_6_ sugar, resulting in 71% carbon conversion efficiency (CCE) of GTs (Table 2G). Roughly 42% of total metabolic CO_2_ was shown to be refixed inside peppermint GTs (Table 2H). In conjunction with its role in supplying precursors for monoterpene biosynthesis, RuBisCO significantly contributed to the fixation of CO_2_, generated in central and monoterpene metabolism. The flux map estimated that RuBisCO assimilated 24% of the CO_2_ released by GT metabolism, thereby strongly contributing to CCE in peppermint GTs (Table 2I). In addition to completely fixing CO_2_ released by high oxPPP flux, RuBisCO assimilated 3% of CO_2_ produced in other processes involved in GT metabolism. Carbon dioxide released by ME was recarboxylated by PEPC in the process of cytosolic PYR production via anaplerotic cycle (Figure 6).

The MFA model integrated the regeneration and consumption of various cofactors (ATP, NADH, and NADPH) to quantify energy and redox-related processes involved in monoterpene biosynthesis. The flux analysis demonstrated that 7.74 mol NADPH was required for 1 mol pulegone biosynthesis (Table 2J) with oxPPP (38%) being the primary supply of NADPH inside GT cells. With respect to NADH metabolism, the TCA cycle represented the main source of NADH production (48%) while glycolysis (17%), monoterpene biosynthesis (13%) and alternate sources (22%) were of minor importance (Table 2K). These results indicated that peppermint GT almost produced sufficient energy for monoterpene production with 51% of NADH being used for ATP production (Table 2L).

## DISCUSSION

### Application of steady state ^13^C-MFA to peppermint GT metabolism

Steady state ^13^C-MFA is a powerful approach for quantifying intracellular metabolic fluxes at the network level (Ratcliffe and Shachar-Hill, 2006; Schwender et al., 2004). Over the last 20 years, steady state flux analysis has been applied in numerous studies to unravel the highly connected pathways of primary metabolism in different plant species (Alonso et al. 2005, 2007, 2010, 2011; Allen et al., 2009, 2012, 2013; Iyer et al., 2008; Kruger et al., 2007; Schwender et al., 2003, 2006; Sriram et al., 2004; Williams et al., 2008). In contrast to that, the more specialized pathways of secondary metabolism are usually analyzed based on kinetic modeling or dynamic labeling studies (Boatright et al., 2004; Heinzle et al., 2007; Matsuda et al., 2005, 2003; Rios-Estepa et al., 2008). Our studied *in vitro* system met the requirements for steady state flux analysis (*i.e*. heterotrophic culture conditions, high ^13^C enrichment, metabolic and isotopic steady state maintained over long time period, and parallel metabolic pathways with carbon backbone rearrangements; Koley et al., 2019). We thus successfully applied steady state ^13^C-MFA to quantify primary and secondary metabolism involved in plant terpenoid production. Important insights in primary and secondary metabolism of non-photosynthetic GTs has been gained in the past years based on genomic and transcriptomic data sets as well as constraint-based modeling results (Ahkami et al., 2015; Croteau et al., 2005; Jin et al., 2014; Johnson et al., 2017; Lange and Srividya, 2019; Lange and Turner, 2013; Tissier, 2012; Zager and Lange, 2018). However, to the best of our knowledge this is the first time that ^13^C-MFA has been applied to provide a quantitative description of GT metabolism involved in terpenoid production.

In peppermint, the complete pathway of monoterpene biosynthesis from imported sugars to monoterpene end products is confined to the non-photosynthetic GT (Gershenzon et al., 1992; McCaskill et al., 1992; McCaskill and Croteau, 1995). Due to their trichome-specific biosynthesis, monoterpenes are well suited to quantify GT metabolism. As shown previously through different studies (e.g. Mendoza-Poudereux et al., 2015), the usage of labeled monoterpenes to quantify terpenoid metabolism does not require trichome isolation. In contrast to that, quantifying GT metabolism by using the isotopic information of intermediates from central and secondary metabolism requires trichome isolation, as those metabolites are not specific to GT and present in different parts of the leaf. Unlike trichome isolation from tomato (Balcke et al., 2017), the isolation of peppermint GTs is not achievable within a few seconds. This procedure takes more than an hour (Gershenzon et al., 1992), during which time the labeling pattern and concentration of metabolites inside GTs would undesirably be renovated at room temperature. Thus, in our study, only GT-specific volatile monoterpenes were investigated to quantify peppermint GT metabolism.

The successful application of steady state MFA depends on selecting appropriate tracers. In many plants of the Lamiaceae family, oligosaccharides of the raffinose family (RFOs) are the most likely primary carbon source imported into GTs (Sprenger and Keller, 2000; Tissier, 2018). As there is no commercial source for ^13^C RFOs (i.e., raffinose), and as RFOs need to be converted into hexoses or hexose phosphates to be further metabolized in the trichome, isotopic GLC was used in our tracer experiments. Based on the current view of RFO catabolism (Madore, 1995; Sprenger and Keller, 2000), the uptake of labeled GLC, its conversion to RFOs for phloem translocation, and its remobilization in the trichome do not involve carbon backbone rearrangement of the GLC molecule, denoting the suitability of GLC as tracer to study GT metabolism. Isotopic GLC was also employed in past investigations to elucidate the isoprenoid biosynthesis in different plants (Lichtenthaler et al., 1997; Mendoza-Poudereux et al., 2015; Schuhr et al., 2003; Skorupinska-Tudek et al., 2008). Early studies in peppermint have also revealed that labeled GLC to be an efficient precursor of monoterpene synthesis (Croteau et al., 1972). Peppermint GTs are typically non-photosynthetic and thus similar to the other peltate trichomes of the Lamiaceae family (Rios-Estepa et al., 2008). While feeding large quantities of GLC has the disadvantage of causing heterotrophic conditions (Shih and Morgan, 2020), this should not affect studies in heterotrophic plant systems such as non-photosynthetic peppermint GTs. The alternative approach of feeding pathway intermediates (such as labeled MVA, H-deoxy-D-xylulose) along with unlabeled GLC was not applied due to difficulties in isotopic steady state establishment and a potential imbalance of natural metabolic pool size which can show biased-favoring of the activity of the respective pathway, thereby causing an elevated interaction between pathways for end-product synthesis. Moreover, isotopic profiling of end product (monoterpene) from one random tracer study might not be sufficient to resolve all fluxes in steady state MFA. Parallel MFA was thus applied in the present study where three positionally different isotopic GLC tracers were selected based on a comprehensive theoretical isotopic tracer analysis within primary and secondary metabolism.

### Significant contribution of the MVA pathway towards monoterpene biosynthesis in peppermint GTs

The present study provided evidence for the contribution of the MVA pathway towards monoterpene production in peppermint GTs. The presence of labeled M+5 and M+6 pulegone from [1,6-^13^C_2_]-GLC demonstrated that the MVA pathway is involved in the biosynthesis of monoterpenes as the production of both mass isotopomers is restricted to the MVA pathway. The formation of these mass isotopomers via the MEP pathway is confined to the provision of M+2 or M+3 PYR resulting from anaplerosis following the repetition of the complete TCA cycle. However, the predicted absence of PYR transport from cytosol to plastid in the present MFA study precludes this possibility (Figure 6). The active contribution of the MVA pathway to the biosynthesis of classical MEP pathway products (e.g. monoterpenes, carotenoids, phytol side chains of chlorophyll) was further proven by the inhibition study. In accordance with these findings, steady state MFA revealed that both the traditional MEP pathway and the alternate MVA pathway contribute to the biosynthesis of monoterpenes. MVA pathway contribution was statistically significant as depicted by the lower bound of the 99.9% CI (6.9%) (Table S5), validating the hypothesis of crosstalk between the two terpene biosynthetic pathways.

Overall, the classical view of peppermint GTs, emphasizes the exclusive role of MEP pathway for monoterpene biosynthesis (Eisenreich et al., 1997). However, the role of the unconventional MVA pathway for monoterpene production was detected in other plant species, such as *Hedera helix* (Piel et al., 1998), *Catharanthus roseus* (Schuhr et al., 2003), *Phaseolus lunatus* (Bartram et al., 2006), *Fragaria sp*. (Hampel et al., 2007), *Gossypium hirsutum* (Opitz et al., 2014) and *Lavandula latifolia* (Mendoza-Poudereux et al., 2015). All these research findings propose an active role of MVA pathway and supports the outcome of the present study. Nevertheless, due to the drawbacks resulting from pathway-inhibitor and pathway-intermediate usage, these studies were restricted to provide a qualitative description of the active existence of the two isoprenoid pathways for the production of certain group of terpenoids. However, our study achieved the quantification of primary and secondary metabolism specific to monoterpene biosynthesis.

### The oxidative bypass of glycolysis fuels MEP pathway

Isotopic tracer analysis and 13C-MFA both suggested that precursors of the classical plastidic MEP pathway were mainly provided by the oxidative bypass of glycolysis. In this oxidative bypass, glycolysis is circumvented by the joint route of oxPPP and RuBisCO.

The oxPPP converts G6P to a pentose phosphate with the generation of NADPH, thus providing reducing equivalents and metabolic intermediates for biosynthetic processes. This path has been expected to be abundant in many oilseeds where NADPH is required for fatty acid biosynthesis (Eastmond and Rawsthorne, 2000; Glas et al., 2012; Kang and Rawsthorne, 1996). In peppermint GTs, phosphogluconate flux (62%) was found to be two times higher than glycolytic fluxes (31%) (Table 2C). Furthermore, transcriptomic and proteomic studies of GTs from spearmint (Jin et al., 2014), peppermint (Ahkami et al., 2015; Zager and Lange, 2018) and tomato (Balcke et al., 2017) also support the indication of an active oxPPP. In contrast to oxPPP, the non-oxidative branch of PPP was shown to carry negligible flux and to act in the direction of pentose phosphate production.

The high metabolic activity of terpene production in GTs requires high amounts of reducing equivalents. For example, the MVA and MEP pathways require 2 and 3 NADPH to produce one IPP (C_5_) from its precursors AcCoA and PYR/GAP, respectively. Additionally, 2 NADPH are needed for the conversion of geranyl pyrophosphate to pulegone. This suggests an elevation of oxPPP flux to support high demand of reducing agents for monoterpene production. Steady state MFA of peppermint GTs suggested oxPPP to be the major source of NADPH generation (Figure 2J). Thus, oxPPP serves a dual function; to provide NADPH as well as precursors for terpene production via the joint route of oxPPP and RuBisCO. Using a genome-scale model of peppermint GT metabolism, Johnson et al. (2017) reported the occurrence of a non-photosynthetic pair of ferredoxin (FD) and ferredoxin-NADP^+^ reductase (FNR) isoforms. By operating in the reverse direction, FNR was predicted to provide reduced FD, thereby recycling NADPH required for monoterpene production. Due to the lack of necessary carbon transfer events, the hypothesis of FNR/FD as an alternate source of NADPH production could not be tested in our ^13^C-MFA study.

Fueled by oxPPP (86%) and non-oxPPP (14%), RuBisCO carboxylates pentose phosphates to triose phosphates, required for monoterpene production. These metabolic characteristics of non-photosynthetic GTs (high RuBisCO and limited non-oxPPP flux) were well supported by ^13^C tracer experiments. In different oil storing green seeds, RuBisCO was shown to act in combination with the non-oxidative branch of PPP (Allen et al., 2009; Schwender et al., 2004). This non-oxidative bypass of glycolysis (RuBisCO shunt) results in an increased carbon conversion efficiency without losing carbon via oxPPP. Unlike non-photosynthetic peppermint GT cells, green embryo of oilseed could accumulate reductant in photosystem-I which perhaps reduces the demand of oxPPP. Additionally, due to significant carbon loss (33%) during synthesis of each C_2_ unit of fatty acid from one C_3_ PYR unit, the biochemical system of oilseed would not prefer further carbon loss via oxPPP.

High carbon use efficiency (more than 70%) in developing seeds of different plant species was observed in previous reports (Alonso et al., 2011; Goffman et al., 2005; Lonien and Schwender, 2009). Our flux map revealed that peppermint GTs are highly efficient in carbon conversion from source to the end product (Table 2G). In our study, RuBisCO was shown to significantly contribute to the fixation of metabolic CO_2_ by assimilating 57% of total refixed CO_2_. Thus, besides its role in supplying precursors for monoterpene production, RuBisCO contributed to the carbon use efficiency in peppermint GTs. In accordance with our findings, transcriptomic studies of GTs from spearmint (Jin et al., 2014) and peppermint (Ahkami et al., 2015) had indicated an active role of RuBisCO. High activity of this enzyme was also reported in tomato and tobacco GTs (Balcke et al., 2017; Cui et al., 2011). Although, GTs of these two Solanaceae species are photosynthetic, the principal role of RuBisCO was reported for reassimilating the internal CO_2_ produced by terpene biosynthesis and other CO_2_ generating metabolic pathways (Balcke et al., 2017; Tissier 2018). A computational analysis of genome-scale models of different GT types predicted RuBisCO to refix metabolically produced CO_2_ in both photosynthetic and non-photosynthetic GTs (Zager and Lange, 2018). This contributes to increased production capacity within these specialized tissues.

To theoretically elucidate the advantage of the oxidative bypass, the non-oxidative bypass and conventional glycolysis with respect to monoterpene production, an elementary flux modes analysis (EFMA) was applied to a simplified version of the ^13^C-MFA model. Details of the EFMA (e.g., model, computation, results) are provided in the supplementary material (File2 and 3). This analysis illustrated three major outcomes. Firstly, the RuBisCO shunt had the highest efficiency in carbon conversion at the cost of requiring the highest amount of NADPH. Secondly, conventional glycolysis had lower CCE, balanced ATP requirements and lower NADPH requirement compared to RuBisCO shunt and thirdly, the oxidative bypass had similar CCE (as the metabolic production of NADPH requires CO_2_ emission), higher ATP demand and lower NADPH requirements compared to the glycolytic route. Taken together, these results pointed to an important role of the oxidative bypass to provide the required NADPH while maintaining balanced CCE for monoterpene production compared to conventional glycolysis and RuBisCO shunt.

### Glycolysis provides precursors for MVA pathway

Flux analysis provided evidence for a significant contribution of glycolytic GAP to supply precursors for the MVA pathway. These MFA-based findings were confirmed by ^13^C tracer analysis of shoot cultures treated with MEP pathway inhibitor. While the MEP pathway starts with the condensation of two trioses (GAP and PYR), three trioses (PYR) are used in the MVA pathway. In case a precursor triose is produced via glycolysis, one extra labeled carbon of [1-^13^C_1_]-GLC is transferred via the MVA pathway (IPP: M+3), compared to MEP pathway (IPP: M+2). (Figure S3). In an alternate route of triose production via the oxPPP and RuBisCO activity, unlabeled IPP is produced from the MVA pathway, contrasting to M+0 and M+1 IPP production via the MEP pathway (Figure 4B.iii). After the partial blockage of the MEP pathway, net ^13^C incorporation was more enriched in pulegone due to a relative increase in MVA pathway involvement. Additionally, labeling phenotype in the FSM-treated culture (i.e. decreased M+0 distribution, and increased distribution in higher mass isotopomers of monoterpene; Figure 5) illustrated the contribution of glycolysis as the principal route of precursor production for the MVA pathway. The present model could not differentiate between cytosolic and plastidic GAP, however, cytosolic source for the MVA pathway could be speculated due to higher influence of sole glycolysis.

Upon glycolysis, AcCoA is metabolized in the mitochondria and subsequently shuttled to the cytosol via the CIT-MAL shuttle to fuel IPP biosynthesis via the MVA pathway. This two-carbon intermediate is impermeable to the membrane and thus synthesized by ATP-citrate lyase in the cytosol (Fatland et al., 2002). Supporting present results of the MFA calculations, transcriptomic data of non-photosynthetic peppermint GT suggested the activation of ATP-citrate lyase as well as other enzymes associated with this shuttle system (Ahkami et al., 2015). Comparative metabolomics, transcriptomics, proteomics, and ^13^C-tracer analysis in photosynthetic GTs of tomato also confirm the role of the CIT-MAL shuttle to supply cytosolic AcCoA required for IPP biosynthesis. Thus, the CIT-MAL shuttle seems to be a very essential component of IPP synthesis towards monoterpene production regardless of the photosynthetic capacity of the trichomes.

To replenish the carbon withdrawn from the TCA cycle by the CIT-MAL shuttle, MAL and PYR are produced via the anaplerotic route of PEPC, MDH and ME in the cytosol. In association with terpene biosynthesis, the anaplerotic reactions have been shown to be active in tomato GTs (Balcke et al., 2017). However, in contrast to this study where plastidic ME was suggested to provide PYR and reducing equivalents for monoterpene biosynthesis in photosynthetic tomato GTs, this enzyme was suggested to be inactive in non-photosynthetic peppermint GT. Further comparative omics study on the non-photosynthetic GTs can help highlight the holistic network of events within these secretory structures.

### Future perspectives of MFA in GTs

The shoot tip culture system in the present study exhibited metabolic and isotopic steady state, which allowed us to investigate the monoterpene biosynthesis in the peppermint GTs using steady state ^13^C-MFA. The combined application of isotopic tracer analysis, ^13^C-MFA and pathway inhibition studies quantified and confirmed the metabolic crosstalk between the two terpene biosynthesis pathways and also the associated primary metabolism. However, a systemic overview of metabolism under natural physiological conditions of the whole plant, might require much tighter control over conditions like light, temperature and humidity. Plants’ sessile nature enables them to evolve or modify their biochemical mechanisms as a rapid response to their ever-changing environment. Light for example is one of the major environmental cues which influences the accumulation of secondary metabolites and could influence the extent of crosstalk between the two pathways. In medicinal plant *Picrorhiza kurroa*, light was shown to play an important role in the expression of both MEP and MVA pathway genes (Kawoosa et al., 2010). Peppermint plants grown at varied light intensities have been reported to accumulate different amounts of terpenes as well as total oil yield (Burbott and Loomis, 1967; Clark and Menary, 1980; Rios-Estepa et al., 2010). Thus, detailed systemic understanding of the metabolic interactions could be improved for example by performing ^13^C-MFA on whole plants using ^13^CO_2_. This could provide even deeper insights into interaction between the MEP and MVA pathway in different GTs regardless of their photosynthetic ability under different environmental conditions.

## MATERIALS AND METHODS

### Chemicals and plant materials

Glucose [C_6_H_12_O_6_] tracers, namely U-^13^C_6_ (99% purity), 1-^13^C_1_ (99% purity), 6-^13^C_1_ (99% purity) and 1,6-^13^C_2_ (99% purity at first position; 97% purity at sixth position) were bought from Campro Scientific GmbH (Berlin, Germany). Fosmidomycin [C_4_H_10_NO_5_P] was purchased from Life Technologies (Carlsbad, USA). Other chemicals and organic solvents were obtained from Sigma-Aldrich (St. Louis, USA) and ROTH (Karlsruhe, Germany), respectively. Peppermint plants (*Mentha × piperita* var. Multimentha) were obtained from Dehner garden center (Halle, Germany). They were maintained in the growth chamber (Microclima MC1000, Snijders Lab, Netherlands) at 25±1 °C, 65% relative humidity and 400 μmol m^−2^ s^−1^ light with 16h photoperiod. Same aged young plants were obtained by vegetative propagation of healthy branches from the purchased plants. About 8-10 cm of the growing branches with leaf buds were dipped into regular tap water within a beaker inside the growth chamber for 2 weeks. During this period, the cutting develops new roots. These cuttings were then planted in the potted moist soil for further growth in the aforementioned climatic conditions.

### Labeling study in shoot tip culture

Shoot tip culture of peppermint plant was performed following the method described by Koley et al. (2019). To perform tracer experiments, these shoot tips were cultivated in MS media at 5 μmol m^−2^ s^−1^ light intensity, which supported maximum ^13^C label enrichment of monoterpene (Koley et al., 2019). In the MS media, GLC was the only carbon source, which varied with the objective of the experiment. From the provision of unlabeled GLC, metabolic steady state of monoterpene production was determined in 13 to 18 days old culture. U-^13^C_6_ GLC were used for establishing isotopic steady state of the growing peppermint shoot tips. For performing MFA, positional labeled GLC (1-^13^C_1_, 6-^13^C_1_ or, 1,6-^13^C_2_) was diluted into the MS media for three independent isotopic studies.

### Inhibition study

Different concentrations of FSM (at 0, 10, 20, 30 and 40 μM) were used for the inhibition of the DXP pathway. Fosmidomycin was added to the MS media, supplemented with unlabeled GLC. First pair of leaves were collected at 15 days old plants and measured for photosynthetic content. Chlorophyll and carotenoid contents were measured in GeneQuant™ 1300 spectrophotometer (Healthcare Bio-Science AB, Uppsala, Sweden), as described in Koley et al. (2019). Labeling pattern and enrichment into pulegone (monoterpene) of 15 days old leaves were also assessed after the partial inhibition of the DXP pathway by 10 μM FSM. As a sole carbon source, 1-^13^C_1_ GLC was supplemented into the inhibitor containing culture media. Isotopic enrichment into monoterpene between control (without FSM treatment) and inhibitor treated leaves were compared for the determination of the active role of the MVA pathway to produce monoterpene.

### Sample collection

Different aged first pair leaves (from 13 days to 18 days old) from the growing shoot tip culture plants were collected for the determination of metabolic steady state. First pair leaves at 14 to 16 days old were collected for isotopic steady state determination, whereas first pair leaves from only 15 days old were collected for other isotopic studies and quantification of photosynthetic pigments. Plant leaf samples were rapidly frozen in liquid nitrogen and stored at −80°C until further investigation.

### Sample extraction and analysis

Frozen leaf samples were pulverised in liquid nitrogen with the help of micro pestles, extracted with hexane and analyzed in a GCMS 2010 gas chromatograph, coupled to GCMS-QP 2010 Plus mass spectrometer (Shimadzu Corporation, Kyoto, Japan) by following the strategies and parameters as detailed by Koley et al. (2019). To assess the metabolic steady state, the relative amount of monoterpenes was measured using nonyl acetate as an internal standard.

### Determination of isotopic enrichment

Average ^13^C enrichment and mass isotopomer distributions (MID) were determined to assess isotopic steady state conditions and to perform MFA. After correcting the raw mass spectrometry data for background noise and natural ^13^C abundance (Fernandez et al., 1996), MIDs and average isotopic enrichment were calculated from four different labeling conditions (1-^13^C_1_ + FSM, 1-^13^C_1_, 6-^13^C_1_ or, 1,6-^13^C_2_ GLC) as follows,

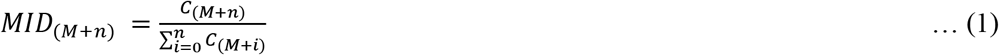

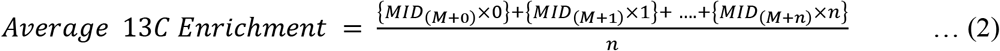

where C and n denote the corrected abundance and number of carbon present in the metabolite, respectively. To determine isotopic steady state and understand the labeling dilution from unlabeled sources (including machine error), average ^13^C enrichment was calculated after further correction for 1% tracer’s impurity from U-^13^C_6_ GLC study.

### Labeling data used for ^13^C-MFA

Steady state ^13^C-MFA was performed based on the isotope labeling measurement of the most abundant mint monoterpenes (i.e. pulegone, isopulegone and menthofuran) from three parallel tracer studies (1-^13^C_1_, 6-^13^C_1_ and 1,6-^13^C_2_). All tracer experiments were performed in triplicates and the average of these were used for flux estimation. In total 9 monoterpene fragments which resulted in 99 MID signals were used for parallel ^13^C-MFA (Table S1, S2 and S3). To account for the effect of natural occurring isotopes, the raw mass spectrometry data was corrected for natural isotope abundance (Fernandez et al., 1996). In addition, the fragments were carefully screened with respect to measurement precision (i.e. natural vs. theoretical isotope abundance ratio). The machine precision (0.03) was used as a threshold value (i.e. min. value of standard deviation) for the standard deviation of all mass isotopomers (Abernathy et al., 2017; Allen et al., 2009).

### Metabolic flux analysis

Isotopic steady state ^13^C-MFA was performed using the MATLAB-based INCA software (Young, 2014), which applies the elementary metabolite unit (EMU) framework (Antoniewicz et al., 2007; Young et al., 2008) for isotopomer analysis. Using a Levenberg-Marquardt optimization algorithm (Dennis and Schnabel, 1983), metabolic fluxes were estimated by least-squares regression of isotope labeling measurements of pulegone, isopulegone and menthofuran from three tracer studies (1-^13^C_1_, 6-^13^C_1_ and 1,6-^13^C_2_ GLC). To perform parallel ^13^C-MFA, the three labeling data sets were simultaneously fit to a single flux model. The metabolic network used for flux calculation is described in the result section and is given in Supplementary Table S4.

All flux determinations were done by normalizing the monoterpene excretion rate to 1 (arbitrary unit). In addition, the upper bound of the ^13^C-GLC uptake was constrained to 3 (arbitrary unit) based on preliminary flux balance analysis computations which predicted a minimal requirement of 1.67 mol GLC for the biosynthesis of 1 mol monoterpenes. As average ^13^C enrichment into monoterpene of 15 days old leaves was 0.5% (Figure 2B), the upper bound of the unlabeled GLC uptake was constrained to 0.01. This value was about 0.5% of labeled GLC uptake. To quantify the energy and redox requirements for monoterpene biosynthesis, the respective reactions for ATP, NADH and NADPH uptake were left unconstrained.

To identify a global best-fit solution, flux calculations were repeated at least 100 times starting from random initial values. The ^13^C-MFA results were subjected to a χ^2^-statistical test to assess the goodness-of-fit between experimental data and model. Given a number of degrees of freedom (DOF) of 56, the estimated fluxes were considered acceptable when the obtained variance-weighted sum of squared residuals (SSR) was below the χ^2^ at a 95% confidence level (χ2 _95%,56_ = 83.3). The statistical uncertainty at 95% CI in the best fit flux values was assessed by evaluating the sensitivity of the SSR to parameter variation (Antoniewicz et al., 2006). Flux precision (i.e. standard error (SE)) was determined as formulated by Antoniewicz et al. (2006) and presented in equation (3).

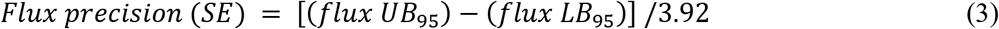

All flux calculations were performed on a 40-core Dell EMC PowerEdge R640 server running Ubuntu LTS 16.04.4 and MATLAB R2017a. Initial flux maps were generated using the Fluxmap add-on (Rohn et al., 2012) of the visualization software VANTED (Junker et al., 2006).

## ACKNOWLEDGEMENTS

The authors thank Dr. Jörg Degenhardt and his scientific group (Managing Director of the Department of Pharmaceutical Biology and Pharmacology, Martin Luther University) for their assistance with GC-MS instrumentation. The authors acknowledge financial support for PhD position from ERASMUS MUNDUS Action 2 program of the European Union under the programme of BRAVE (Grant Agreement Number 2013-2536/001-001).

## AUTHOR CONTRIBUTIONS

SK and BJ designed the research. SK performed the wet-lab experiments and EGB performed the mathematical modelling. SK, EGB, MLR and BJ analyzed the data and wrote the manuscript. All authors read and approved the manuscript.

